# The thermoception task: a thermal-imaging based procedure for measuring awareness of changes in peripheral body temperature

**DOI:** 10.1101/2022.09.07.506983

**Authors:** Alisha Vabba, Maria Serena Panasiti, Marina Scattolin, Marco Spitaleri, Giuseppina Porciello, Salvatore Maria Aglioti

## Abstract

Although thermal body signals provide crucial information about the state of an organism, and changes in body temperature may be a sign of affective states (e.g., stress, pain, sexual arousal), research on thermal awareness is limited. Here we developed a task measuring awareness of changes in peripheral body temperature (*thermal interoception*) and compared it to the classical heartbeat counting task (*cardiac interoception*). With an infrared lightbulb we delivered stimuli of different temperature intensities to the right hand of 31 healthy participants. *Thermal interoceptive accuracy*, i.e., the difference between participants’ real and perceived change in hand temperature, showed good inter-individual variability. We found that thermal interoception did not correlate with (and was generally higher than) cardiac interoception, suggesting that different interceptive channels provide separate contributions to awareness of bodily states. Moreover, the results hint at the great salience of thermal signals and the need for thermoregulation in day-to-day life. Finally, thermal interoceptive accuracy was associated with self-reported awareness of body temperature changes, and with the ability to regulate distress by focusing on body sensations. Our task has the potential to significantly increase current knowledge about the role of interoception in cognition and behavior, particularly in social and emotional contexts.

## Introduction

Recent years have shown increasing interest in the role that awareness of the physiological state of the body, or interoception ^1,2^, plays in emotion ^for recent reviews see 3–5^, psychopathology ^for a review see 6^, and social cognition and behaviour, such as empathy and perspective taking ^7–11^, social disposition ^12^, altruistic behaviour ^13^, moral identity ^14^, and moral decision-making ^15^. Research on interoception has focused mainly on the cardiac domain, because heartbeats are distinct events that can be easily measured ^16^ and because of the important role that heart-brain interactions seem to play in emotional processing ^17^. However, the tasks employed in the assessment of cardiac interoception and heartbeat estimation have recently come under severe controversy ^18,19^, which raises questions about their psychometric validity. Indeed, performance in cardioceptive tasks was found to be influenced by time estimation (mentally counting the time of different trials) and prior beliefs concerning one’s heart rate ^20^, even when these beliefs are induced through false feedback prior to performing the task ^21^. Additionally, Windmann and colleagues found that the number of heartbeats reported by participants did not change even when real heartbeats were manipulated using cardiac pacemakers ^22^.

Moreover, it is important to consider that, the construct of interoception refers not only to cardiac signals, but to a larger number of sensations including hunger, thirst, dyspnoea, sexual arousal, cooling or warming, bladder distension, and stomach sensations ^2^. Tellingly, growing evidence suggests that cardiac interoception cannot be taken for a proxy of interoceptive accuracy across all sensory channels. For example, studies have found that cardiac interoception does not correlate positively with respiratory interoception ^23^, gastric interoception ^24,25^, pain ^25,26^, nor with skin-mediated thermal interoception, or sensual touch ^26^. A limitation of some existing measures assessing non-cardiac interoception is that they are often based on invasive procedures, such as air load judgements ^27^ and gastric ^28,29^ and bladder ^30^ balloon distention procedures. The limitations of existing measures and the notion that cardiac interoception cannot be generalised to all interoceptive domains call for the need to develop new and non-invasive protocols tapping into different interoceptive channels.

To our knowledge, one aspect of interoception that has so far received less attention, regardless of its fundamental role in survival, is the perception of body temperature, which can be mediated by the external environment but also arise inside the body, as happens with fever. Indeed, the survival of most species - humans included - depends on an optimal body temperature ^31^, and human beings make use of complex autonomic and behavioural means to survive and thrive while exposed to a wide range of environmental temperatures. In fact, dysregulations in body temperature – particularly in these global warming times, may have serious, dramatic effects: for example, the mortality rate among people with compromised thermoregulation capacities, like the elderly ^32^, increases during extremely hot or cold seasons ^33^. Thermosensation, i.e., the perception of (peripheral and core) body temperature is considered an interoceptive modality as it provides important information about the body’s physiological state and shares similar pathways to other interoceptive modalities (i.e., the spinal laminar 1 spinothalamic tract which carries information relevant to homeostatic control and physiological needs) ^2^.

Recent evidence showed that visuo-thermal incongruencies reduced the illusory experience of owing a rubber hand and gave rise to visuo-thermal illusion effects, suggesting that thermal information is integrated with other bodily signals and crucially contributes to our sense of body ownership ^34^. Furthermore, our awareness of body temperature can affect our reliance on and employment of strategies aimed at regulating the body temperature i.e., behavioural thermoregulation ^35,36^ and may be important in emotional and stressful contexts where body temperature can reveal changes in autonomic function. For example, increases in facial temperature have been associated with a wide variety of affective states such as emotional ^37–39^, anxiety ^40^ and sexual arousal ^41^, stress and pain ^41^. In social scenarios, increases in facial temperature have been highlighted in response to social contact ^42^ and social ostracism ^43,44^, during dishonest behaviour ^45,46^ and in-group outgroup categorization ^47^.

Thus, tasks aimed at quantifying thermal interoception, can be an important instrumental tool in the experimental study of interoception and body awareness, with relevant applications for exploring the role of bodily changes during social and emotional contexts. To our knowledge, only a few measures of awareness of temperature changes are available. Heldestad et al., (2010) developed a measure of thermosensitivity using the method of limits ^MLI; 49^ to deliver cold and warm stimuli with a thermode placed on the lower arm, that started at 32°C and gradually increased/decreased. Participants had to report when they perceived a thermal sensation to derive a threshold of thermal perception. Quite recently, a thermal matching task has been developed ^26,34^ in which participants’ forearms or palms were stroked with a thermode at a reference temperature of 30°, 32°, or 34°C. Next, the experimenter touched the participant with different temperatures starting from ± 8°C of the reference temperature and gradually increasing/decreasing. Participants indicated when they thought the reference temperature was reached. However, all the above tasks used tactile stimulations to deliver thermal stimuli and required participants to give judgements about an external object and not the temperature of their own arm/hand.

The aim of the present study was to develop a task that measured awareness of changes in peripheral body temperature without requiring the tactile modality nor the estimation of the temperature of an external object. During this task, we used infrared radial thermal stimulation of different temperature intensities to induce changes in temperature to the palm of participants’ right hand. We then asked participants to give judgements related to the perceived temperature of their hand, rather than the temperature of an external object. This way the task was expected to reduce tactile related confounds, and to measure interoceptive accuracy as an index of awareness of the physiological state of the body included motivated signals from the skin surface ^1,2^. We also asked participants to rate their perceived level of confidence in task performance as a proxy to measure meta-awareness. Participants also underwent the heartbeat counting task and completed a series of self-reported measures of *thermosensitivity* i.e., selected sub-scales of the Experienced Temperature Sensitivity and Regulation Scale, ETSRS ^50^ and the Social Thermoregulation and Risk Avoidance Questionnaire, STRAQ-1 ^51^ and measures of *interoceptive sensibility* i.e., the Multidimensional Assessment of Interoceptive Awareness, MAIA-2 ^52^ and the Body Perception Questionnaire, BPQ-SF ^53^.

We created indices of thermal interoceptive accuracy (the capacity to accurately detect changes in hand temperature) and awareness (i.e., correspondence between accuracy and confidence in task performance) and used them to compare the participants’ performance in the thermal task and the heartbeat counting task. We predicted a dissociation between thermal interoceptive accuracy and awareness, mirroring the multi-dimensionality of cardiac interoception ^54^. Furthermore, with respect our main hypothesis namely testing whether interoceptive channels correlate between each other, based on literature suggesting a dissociation between interoceptive sub-modalities ^23–26^, we would expect that thermal interoceptive accuracy would be unrelated to cardiac interoceptive accuracy, whereas based on literature showing a correlation between certain sub-modalities ^24^ we would expect that the two dimensions may be related. Finally, we set out to validate the present task by testing whether performance in the thermoception task could be predicted by questionnaires assessing thermosensitivity (ETSRS and STRAQ-1 scores) and general interoceptive sensibility (the MAIA-2 and BPQ-SF scores).

## Methods

### Participants

A power analysis (*power* = 0.8; *α* = 0.05; effect size decided on the basis on the average correlations (*r* = 0.49) found by previous studies comparing tasks across different interoceptive modalities (Van Dyck et al., 2016; Whitehead & Drescher, 1980; Herbert et al., 2012)) using the *pwr* package for R, was used to estimate the sample size need for running a correlation analysis between the performance in the thermoception task with performance in a task of cardiac interoception (the Heartbeat Counting Task, Schandry, 1981). According to the power analysis a sample of 30 participants would be sufficient to detect a significant correlation between the tasks. A total of 47 participants took part in the study. However, the experimental protocol required strict control of the environmental temperature and, due to Covid-19 regulations related to ventilation of the testing room, these conditions were violated in a great number of cases, leading to the elimination of the responses of 16 participants. The final sample consisted of 31 healthy volunteers (16 males, age [*mean* = 27.21, *standard deviation* = 4.71]). The environmental temperature of the testing room was continuously checked by means of an external thermometer and we managed to keep it constant throughout the experiment. The study was approved by the Ethics Committee of the Santa Lucia Hospital I.R.C.C.S. and was in accordance with the 1964 declaration of Helsinki.

### General procedure

Upon arrival at the laboratory, participants were informed they were taking part in a study aimed at validating a task for the assessment of awareness of changes in body temperature. During the first 30 minutes, we asked participants to comfortably sit on a chair while they acclimatised to the environment and to the room temperature. During this time, they were asked to read and sign the informed consent. Then they performed the thermoception task and the heartbeat counting task, in counterbalanced order (see below for a detailed description of the two tasks). A series of self-reported measures were completed by the participants at home, using the *Survey Monkey* platform (see below for the detailed description of the selected questionnaires).

### Behavioural measures

#### The thermoception task

*Figure 1*. Panel A. (see below) depicts the task set-up. For the entire duration of the thermoception task, participants were seated at a desk. During each trial, the participants placed their right hand on a hand rest, which was secured to the desk and covered in thermal foil. Opposite the hand rest, at 15 cm, was an infrared light bulb (radiant heating; maximum power: 250 watt; voltage: 240 Volt; dimmable; dimensions: 12 × 12 × 17.5 cm; weight: 160 g), which delivered stimulations of different thermal intensities. Specifically, the experimenter used a dimmer to set the lightbulb power at 0%, 25%, 50%, 75%, and 100%. A FLIR A655sc thermal camera (FLIR Systems, AB, Sweden) was positioned next to the lightbulb, so to frame the right hand of participants and record its temperature throughout each trial. To bring the temperature of the stimulated hand back to a baseline value before the following trial, participants immersed their right hand in a thermostatic water bath (FALC Instruments model WB-MD15) placed to their right. On their left, participants had a computer screen and a mouse, which they used to provide their estimations of temperature change (see below for the detailed description of the procedure). To avoid responses from being biased by visual cues, a shield was placed in front of the participant so that the light bulb was covered from view.

**Figure 1.**
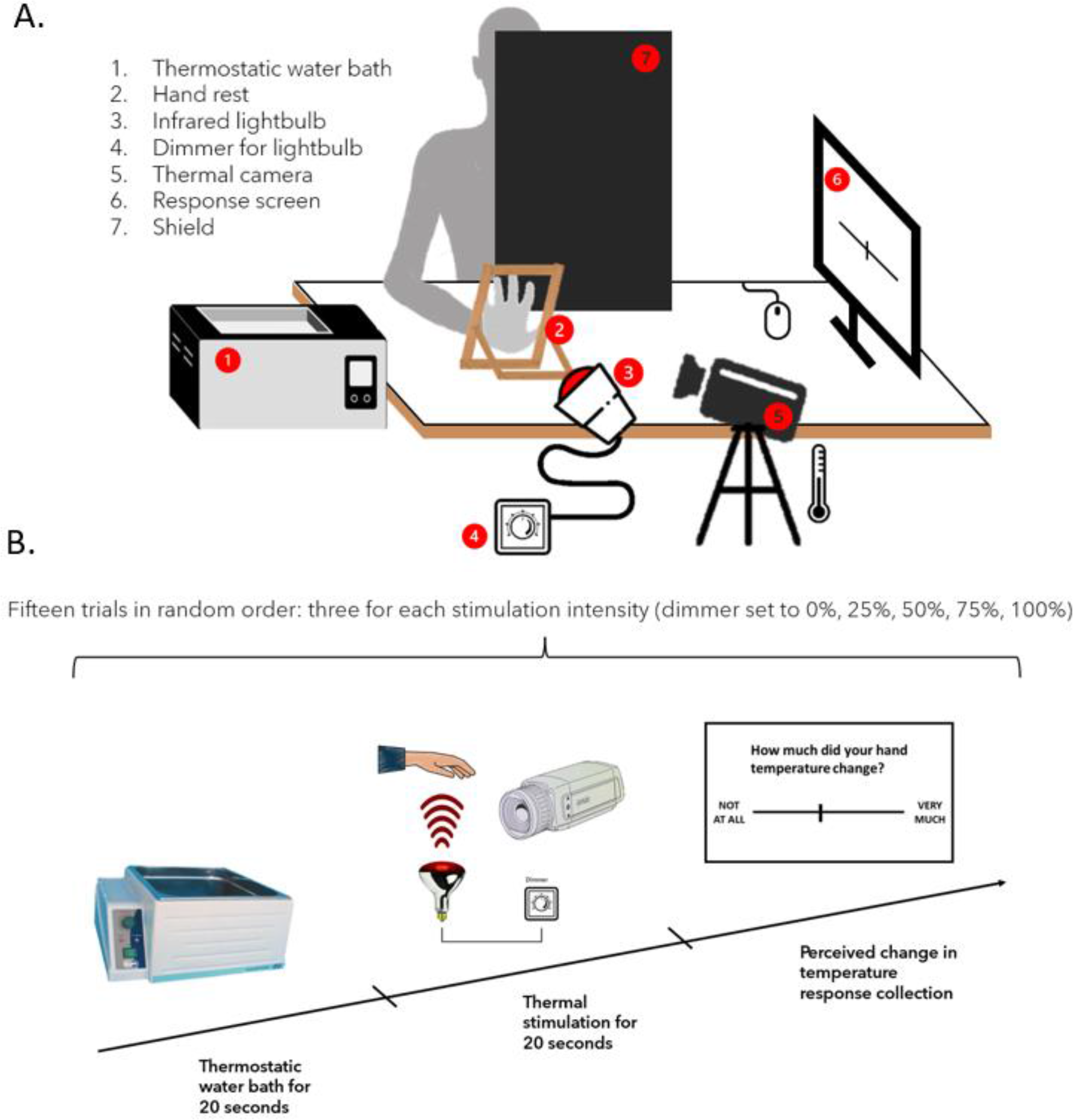
Experimental set-up and design. Panel A. The thermoception task set-up: Participants are seated at a desk with a rest for their right hand. On their right is a thermostatic water bath, and on their left are a screen and a mouse for instructions and response collection. An infrared light bulb is attached to the desk at a 15 cm distance from the hand. A shield covers the lightbulb from view, to avoid response bias due to visual cues. A thermal camera is used to record real changes in hand temperature during the task. Panel B. Illustration of a typical trial of the thermoception task. The task consists of 15 trials, three per each temperature intensity (0%, 25%, 50%, 75%, 100%). In each trial: 1) the participants insert their right hand for 20 seconds in the thermostatic water bath set at 31.5 °C; 2) participants dry their hand, place it on the stand and receive the 20 second thermal stimulation, delimited by two acoustic tones; 3) participants report the perceived change in temperature from before to after the stimulation on a VAS ranging from 0 = “Not at all” to 100 = “Very much”.

The experiment (see below *Figure 1*. Panel B) consisted of 15 trials (three per intensity of stimulation presented in randomised order), in which participants received thermal stimulations to the palm of their right hand. Prior to this, participants completed three practice trials: these served as a way for participants to familiarize with the task procedure and with the highest stimulation intensity. As such, practice trials were not considered in the statistical analysis.

Each experimental trial was structured in the following three steps:

1. At the beginning of each trial, for 20 seconds, participants immersed their right hand in the thermostatic water bath. Water temperature was set to 31.5 °C, so to bring the hand temperature to a comparable and controlled baseline value before the following stimulation. During these 20 seconds, and in accordance with the randomized order generated by E-Prime, the experimenter set the dimmer to the correct intensity. The dimmer was hidden from participants’ sight for the entire duration of the study.
2. When prompted by the experimenter, the participants quickly and thoroughly wiped their hand using a paper towel and then placed it back on the hand rest. Then, they were instructed to close their eyes and concentrate on the current temperature of their hand. In fact, participants’ task was to compare the temperature of their hand at two time points, that were signalled by two acoustic tones. Once participants were ready, the experimenter launched the first tone. After two seconds the light bulb was turned on and switched off after 20 seconds, i.e., when a second acoustic tone was delivered. Participants were then instructed to open their eyes and removed their hand from the stand.
3. When, the thermal stimulation ended, participants read the following question “*How much did the temperature of your hand change from the first to the second acoustic tone*?” on the screen in front of them. Responses were provided with a left mouse click, along a visuo-analogue scale (*perceived temperature change VAS*), which ranged from *0 = “Not at all”* to *100= “Very much”*.

At the end of the thermoception task, participants rated how accurate they believed their performance in the whole task to be. They did so on a second, visuo-analogue scale (*confidence rating VAS*), that in this case ranged from 0 = “*Not accurate at all”* to *100 = “Completely accurate”*. Instructions, trial order, thermal camera triggering, and response collection were handled through E-Prime 2 software (Psychology Software Tools, Pittsburgh, PA).

The real change in the participants’ hand temperature was recorded via a thermal camera using the software FLIR Research IR Max 4. The camera was triggered to start and stop recording respectively when the first and second acoustic tone were delivered, thus producing 15 files per participant.

#### The heartbeat counting task

During the heartbeat counting task ^HCT; 55^ participants are asked to mentally count the number of heartbeats they perceive in the absence of physical cues (e.g., taking their pulse) during four intervals of different duration (i.e., 25, 35, 45, and 100 seconds). The order of the time intervals is randomized, and each is delimited by two auditory tones that the participants hear through a headset. At the end of each trial, participants enter the number of heartbeats they have felt, using a computer keyboard. In the present study, participants were explicitly asked not to *estimate* the number of heartbeats they believed occurred during the trial, but to only consider heartbeats they truly perceived. At the end of the experiment, participants rated their perceived level of accuracy in the task, using a visuo-analogue scale (VAS) ranging from 0 = “*Not accurate at all”* to *100 = “Completely accurate”*. Cardiac signals were recorded throughout the task using a physiological recording system (ADInstruments PowerLab and BIOPAC). Instructions, trial order, and response collection were handled by E-Prime 2 software (Psychology Software Tools, Pittsburgh, PA).

### Self-report measures

The **Experienced Temperature Sensitivity and Regulation Scale** ^ETSRS; 50^ is a questionnaire designed to assess perceived thermosensitivity and autonomic or behavioural thermoregulatory activity. We selected and administered two sub-scales: the *Heat-induced warming* subscale is composed of 5 items and measures participants’ subjective perception of how quickly or intensely different body parts, like, for example, hands, feet and torso, become warm when exposed to warm environments (e.g., *“Compared to others, a warm environment gives me warm feet*”) and the 7-item, *Heat perception* subscale measuring the self-reported tendency to feel warm in different environments and/or situations e.g., indoors, outdoors, in bed, when concentrating, when watching TV or reading (e.g., “*Compared to others, I experience heat at home*”). For both subscales, participants judge how much each sentence is true for them compared to other people, using a 6-point Likert scale ranging from *1= “Much less”* to *7 = “Much more”*.

The **Social Thermoregulation and Risk Avoidance Questionnaire** ^STRAQ-1; 51^ is a survey developed to measure the importance of thermal cues and thermoregulatory biological drives in the formation of attachment and social relations (i.e., seeking physical contact and closeness to other people). We selected and administered three sub-scales: the *High temperature Sensitivity* 7-items subscale, that measures how sensitive to and bothered by high temperatures participants are (e.g., “*I can’t focus when it is too hot*”); the *Solitary Thermoregulation* 8-items subscale that estimates the tendency to use behavioural thermoregulatory actions to maintain thermoneutrality (e.g. “*When it is cold, I wear more clothing than others*”); the *Social Thermoregulation* 5-items subscale, that reflects the tendency to be physically close and to physically warm up with other people (e.g., “*I like to warm up my hands and feet by touching someone I am close to*”).

The **Multidimensional Assessment of Interoceptive Awareness** ^MAIA-2; 56^ is a 37-item survey measuring interoceptive sensitivity and it is composed of 8 sub-scales, specifically *Noticing* (4 items, e.g. “*When I am tense, I notice where the tension is located in my body*”), which measures awareness of uncomfortable, comfortable, and neutral body sensations; *Not-Distracting* (6 items, e.g. “*I distract myself from sensations of discomfort*”), assessing the tendency not to ignore or distract oneself from sensations of pain or discomfort; *Not-Worrying* (5 items, e.g. “*I start to worry that something is wrong if I feel any discomfort*”), measuring the tendency not to worry or experience emotional distress with sensations of pain or discomfort; *Attention Regulation* (7 items, e.g. “*I can pay attention to my breath without being distracted by things happening around me*”), which assesses the ability to sustain and control attention to body sensations; *Emotional Awareness* (5 items, e.g. “*I notice how my body changes when I am angry*”) measures awareness of the connection between body sensations and emotional items; *Self-Regulation* (4 items, e.g. “*When I feel overwhelmed, I can find a calm place inside*”), measuring the ability to regulate distress by attention to body sensations; *Body Listening* (3 items, e.g. “*I listen for information from my body about my emotional state*”) assesses active listening to the body for insight; *Trusting* (3 items, e.g. “*I am at home in my body*”) measures the experience of one’s body as safe and trustworthy. Responses are collected on a 6-point Likert scale ranging from *0 = “Never”* to *5 = “Always”*.

**The Body Perception Questionnaire-Short Form** (*BPQ-SF*; Cabrera et al., 2018) is a 22-item questionnaire assessing self-reported body awareness and autonomic reactivity. It measures the tendency to be aware of different body sensations and processes, such as bloating, stomach sensations, sweating, tremor, facial temperature (e.g., “*My heart often beats irregularly*”, “*I feel shortness of breath*”). Participants report how often they are aware of specific sensations using a 5-point Likert scale ranging from 1= “*Never”* to 5= “*Always*”.

### Analytic plan

#### Measuring real and perceived hand temperature changes

For every participant, each of the 15 thermal video files recorded by the thermal camera was visually inspected to make sure participants were positioned correctly and did not move during the trial. An area of interest (AOI) was designed to comprise the palm of the participant’s hand with the software FLIR Research IR Max 4, using the base points of the palm and of each finger as reference points (see *Figure 2* for the AOI and a photographic example of thermal recordings across all stimulation intensities). Being directly affected by the stimulation, the defined area may be optimal for detecting thermal changes.

**Figure 2.**
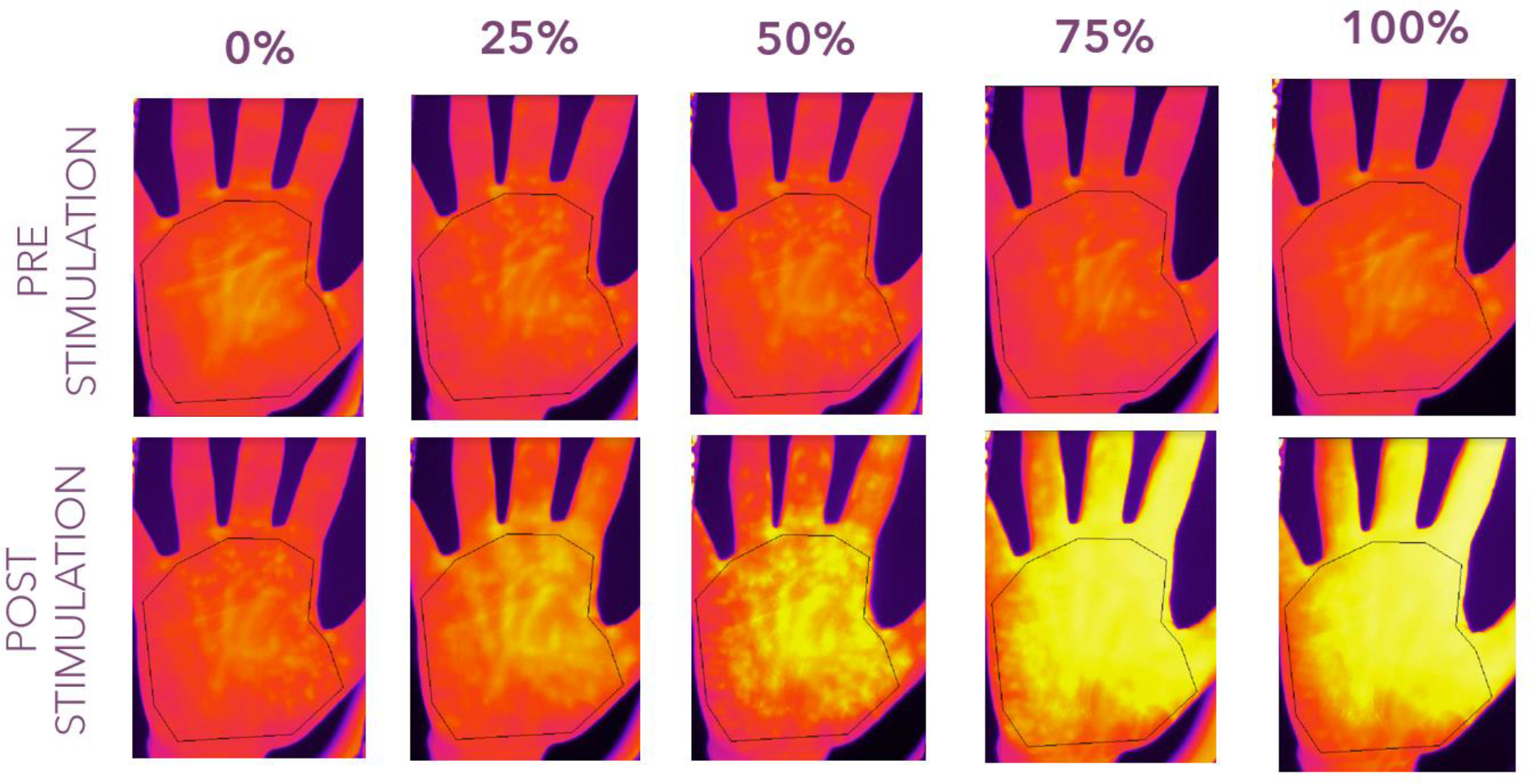
Thermal images of a representative participant’s hand. Thermal images were recorded via the thermal camera FLIR A655sc and FLIR Research IR Max software. The images were taken at baseline (two seconds before stimulation) and at the end of the 20-second stimulation for each of the five intensity conditions (0%, 25%, 50%, 75%, 100%).

For each point in time, the average temperature (in degrees Celsius) within the area of interest was outputted into a single csv file. The first and last entries in the csv file were used respectively as a measure of hand temperature before and at the end of the thermal stimulation. We then calculated the difference in temperature between the two time points, i.e., the *real hand temperature change* for each trial.

The *perceived hand temperature change* for each trial was the score in the perceived temperature change VAS.

#### Effect of stimulation intensity on real and perceived hand temperature change and thermal interoceptive accuracy

To test whether different stimulation intensities induced different *real* and *perceived hand temperature changes*, and different scores in *thermal interoception accuracy* (see next section *Measuring thermal interoceptive accuracy and awareness*) we performed multilevel mixed model analyses using the R package *lme4 v 1.1-23*. Multilevel mixed model analysis was used to account for missing data due to technical issues with some of the trials. We constructed three separate models with the dependent variable respectively set to 1) *real hand temperature change*, 2) *perceived hand temperature change* and 3) *thermal interoceptive accuracy*, with stimulation intensity (0%, 25%, 50%, 75%, 100%) as the main within-subject predictor. Each trial of each participant was considered as a separate observation, therefore we entered 15 observations per participant. To deal with the non-independence of the dataset, the random intercept was set on Participant. We conducted a *Type III Wald Anova* using the *car* package for R (R Studio version 4.0.2). Post-hoc comparisons were performed with the *R emmeans* package and with Bonferroni correction for multiple comparisons.

#### Measuring thermal interoceptive accuracy and awareness

We first transformed the *real hand temperature change* for each trial into a value between 0 and 100 (*standardized hand temperature change*), to better compare it with the score of *perceived temperature change* which was rated on a 0-100 VAS. For each participant, we set the larger temperature change across all 15 trials to be the upper end of the range, while 0 indicated no change in hand temperature. We then transformed the real temperature change in each remaining trial according to the following formula:

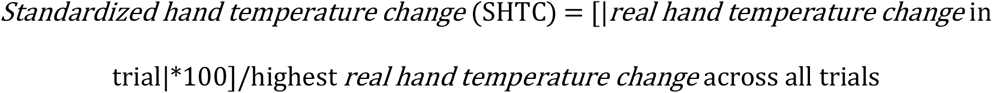

SHTC values of 0 and 100 indicate, respectively, no change and the largest possible temperature change.

To have a unique measure of *thermal interoceptive accuracy* for each participant we used the following formula:

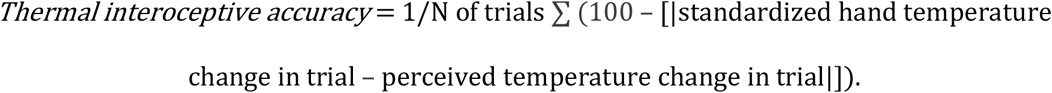

Values of 0 and 100 indicate worst and best thermal interoceptive accuracy, respectively.

*Thermal interoceptive awareness* was then calculated using the following formula:

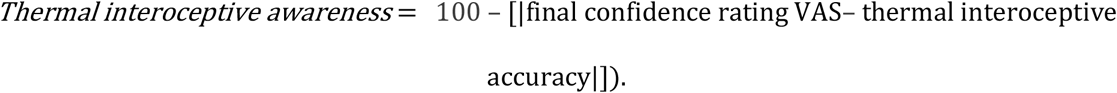

Values of 0 and 100 indicate, respectively, no awareness and perfect awareness.

#### Measuring cardiac interoceptive accuracy and awareness

Cardiac interoceptive accuracy was calculated as the difference of perceived to real heartbeats averaged across all trials, using the following formula:

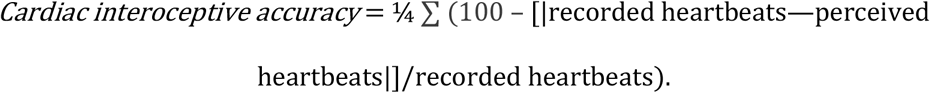

Scores closer to 100 indicate higher accuracy.

Cardiac interoceptive awareness was calculated as the difference between confidence judgements (scores in the final confidence rating VAS) measured on the overall performance in the HCT and the cardiac interoceptive accuracy score.

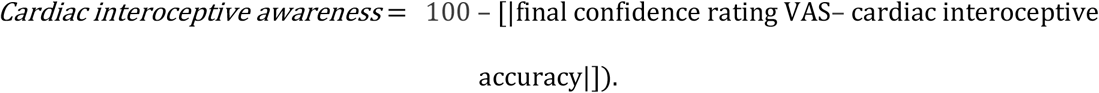

Values of 0 and 100 indicate, respectively, no awareness and perfect awareness.

#### Comparison and correlation analysis between thermal and cardiac interoception

To test the difference between thermal and cardiac interoceptive accuracy and awareness we performed t-tests. To test covariation between thermal interoceptive accuracy and awareness and cardiac interoceptive accuracy and awareness, we performed Bonferroni-corrected Pearson’s correlations and, where required, we calculated effect sizes and Bayes factor, using JASP software (JASP v.0.14.1). Bonferroni cutoff for significant correlations was determined by dividing the conventional *p*= .05 cutoff by the total number of correlations we computed (i.e., 4), which thus corresponded to *p* = .012. In all cases, as thermal interoceptive accuracy did not vary across active stimulations (25%, 50%, 75%, and 100%), which all differed from the baseline stimulation (0%), thermal interoceptive accuracy and awareness were calculated based on the average of the 13 active stimulation intensity trials (25%, 50%, 75%, 100%) excluding the baseline trials (0 %).

#### Exploratory regression analyses between thermal interoception and subjective measures of thermosensitivity and interoceptive sensibility

We performed linear regression analyses (linear models, LM) using the R package *lme4 v 1-1-23* to investigate whether thermal interoceptive accuracy was predicted by subjective thermosensitivity and general interoceptive sensibility. Considering that thermal interoceptive accuracy did not vary across active stimulations (25%, 50%, 75%, and 100%), which all differed from the baseline stimulation (0%), we only included active stimulations in the models. Both models had thermal interoceptive accuracy (for active stimulations) as the dependent variable. The model assessing the relation between thermal interoceptive accuracy and thermal sensitivity corresponded to the following:

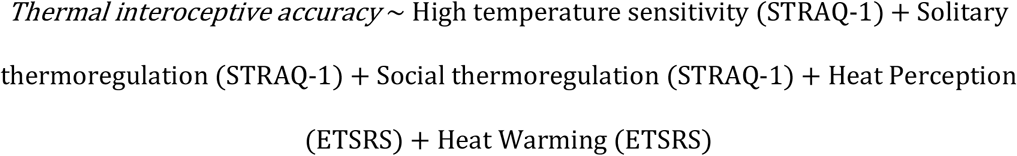

The model for investigating the role of interoceptive sensibility (MAIA-2 subscales and the BPQ-short form) in thermal accuracy was coded as follows:

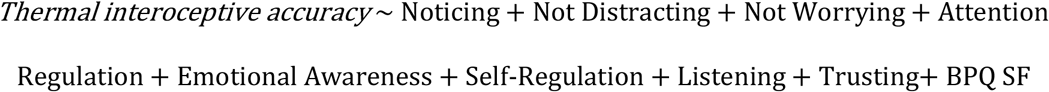

We conducted Type III Wald Anovas using the car package for R.

## Results

### Environmental temperature

Average room temperature at the beginning of the experiment was 25.69 °C (*standard deviation* = 1.17). The change in room temperature from the beginning to the end of the experiment was of 0.645 °C on average (*standard deviation* = 0.65). The average temperature of the thermostatic water bath at the beginning of the experiment was 31.53 °C (*standard deviation* = 0.09) and remained constant throughout the experiment for all participants.

### Effect of stimulation intensity on real and perceived change in hand temperature and on thermal interoceptive accuracy

#### Real hand temperature change

Mean baseline hand temperature (i.e., taken at the first acoustic tone corresponding to the first real thermal frame) for all participants and stimulation types was 32.39 °C (*standard deviation* =1.24). There was a significant main effect of stimulation intensity on the *real hand temperature change* (*χ*^2^ = 4027.60, *p* < .001). Post-hoc analysis revealed that the temperature change was significantly higher for all stimulation intensities (25%, 50%, 75%, and 100%) compared to the baseline, 0% stimulation intensity (smallest *b* = −0.65, smallest *t* = −14.72, all *p*-values < .001). Furthermore, *real hand temperature change* was significantly higher for the 50% compared to the 25% stimulation intensity (*b* = −.841, *t* = −18.99, p< .001), for the 75% compared to the 50% stimulation intensity (*b* = −0.62, *t* = −14.01, p< .001), and for the 100% compared to the 75% stimulation intensity (*b* = −0.24, *t* = −5.48, *p*< .001). (*Figure 3*, Panel A.)

**Figure 3.**
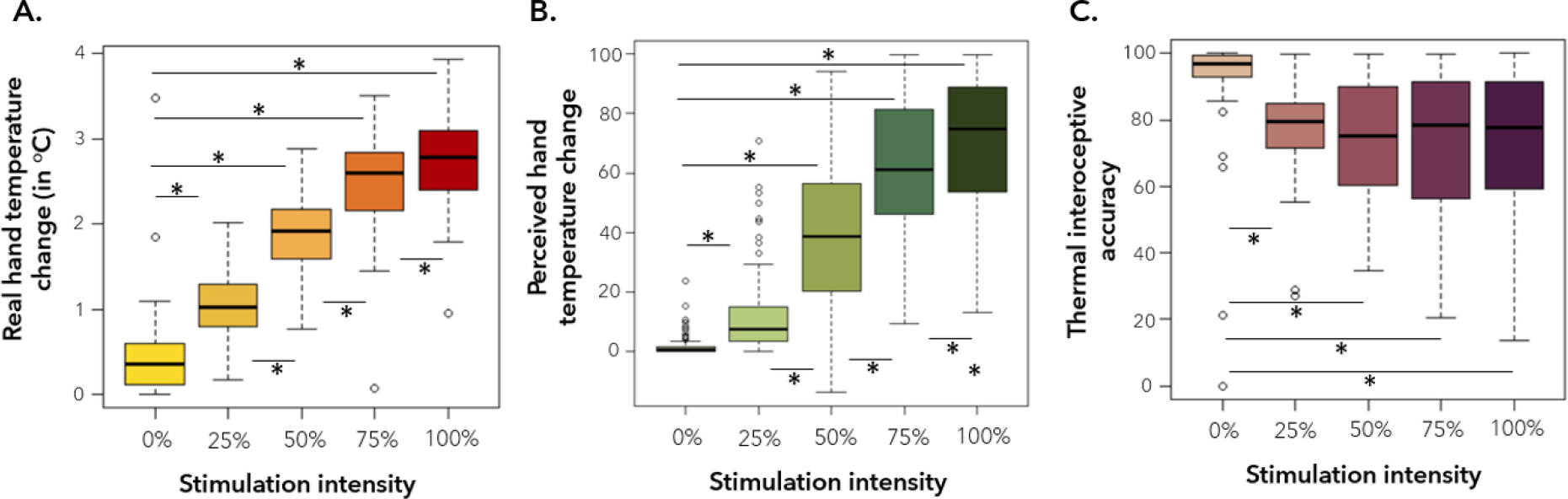
Box plots displaying differences in real and perceived change in hand temperature and interoceptive accuracy based on stimulation intensity. Values for each stimulation intensity are reported (0%, 25%, 50%, 75% and 100%). Panel A represents differences in real change in hand temperature from before to after the stimulation, recorded by a thermal camera in °C. Panel B shows differences in average perceived hand temperature (average of VAS scores). Panel C shows differences in thermal interoceptive accuracy. * indicates *p* < .001

#### Perceived hand temperature change

There was a significant main effect of stimulation intensity on *perceived hand temperature change* (*χ*^2^ = 1635.32, *p* < .001). Post-hoc analysis revealed that the perceived temperature change was significantly higher for all stimulation conditions (25%, 50%, 75%, and 100%) compared to the 0% stimulation intensity (smallest *b* = −10.57, smallest *t* = −5.19, all *p-value*s < .001). Furthermore, *perceived hand temperature change* was significantly higher for the 50% compared to the 25% stimulation intensity (*b* = −27.74, *t* = −13.59, p< .001), for the 75% compared to the 50% stimulation intensity (*b* = −21.99, *t* = −10.77, p< .001), and for the 100% compared to the 75% stimulation intensity (*b* = −8.36, *t* = −4.101, p< .001) (*Figure 3*, Panel B.)

#### Change in thermal interoceptive accuracy

There was a significant main effect of stimulation intensity on thermal interoceptive accuracy (*χ*^2^ = 99.69, *p*< .001). Post-hoc analysis revealed that participants were more accurate at baseline compared to all other stimulations (smallest *b* = 14.17, smallest *t* = 7.42, all *p-values*< .001). There was no significant difference in the capacity of participants to detect temperature changes across the other stimulation intensities (all *p-values* > .05). (*Figure 3*, Panel C.).

### Cardiac and thermal interoception

#### Characterization of thermal interoception

Considering that thermal interoceptive accuracy did not vary across active stimulations (25%, 50%, 75%, and 100%), which all differed from the baseline stimulation (0%) we used only the active stimulations to compute a general measure of *thermal interoceptive accuracy*. Our measure of thermal interoceptive accuracy was generally quite high (*mean*= 73.07, *standard deviation* =16.45), presented high inter-individual variability (ranging from 41.44 to 91.59), and was internally consistent across different types of stimulation intensity (*α*= .820). Thermal interoceptive awareness was also high (*mean* = 84.96, *standard deviation* =13.49), ranging from 53.21 to 99.36.

#### Characterization of cardiac interoception

Cardiac interoceptive accuracy (*mean* = 47.81, *standard deviation* = 23.45), was also internally consistent (*α*= .928) and presented high inter-individual variability, ranging from 0 to 97.8. Cardiac interoceptive awareness was high (*mean* = 0.88, *standard deviation* = 0.96), varying from 33.27 to 97.34.

#### Comparison and covariation between thermal and cardiac interoception

The distribution of cardiac and thermal interoception scores concerning accuracy and meta-awareness are represented in *Figure 4*.

**Figure 4.**
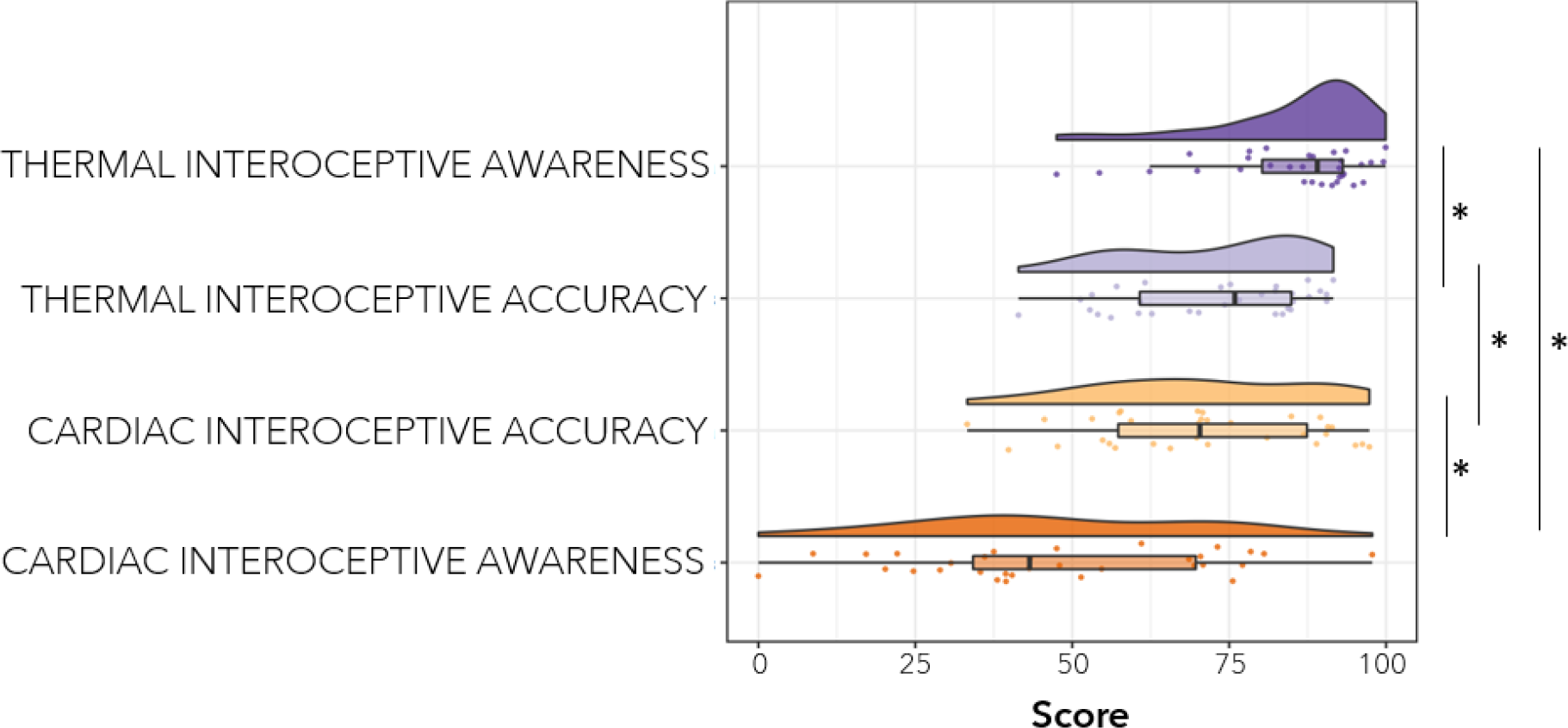
Raincloud plots displaying the distribution of raw data, box plots, and split violin plots for thermal and cardiac interoception (accuracy and awareness scores). Values closer to 100 indicate higher scores. *indicates a significant difference in scores at p< .001

*Thermal interoceptive accuracy* scores were significantly higher than *cardiac interoceptive accuracy* (*t* = −5.17, *p* < .001). *Thermal interoceptive awareness* scores were also significantly higher than *cardiac interoceptive awareness* (t= −3.85, p < .001).

*Thermal interoceptive accuracy* did not correlate with *thermal interoceptive meta-awareness* (*r* = 0.30, *p* = .090, *BF* = 0.871), and similarly *cardiac interoceptive accuracy* did not correlate with *cardiac interoceptive awareness* (*r* = − 0.25, *p*= .165, *BF* = 0.554). For both correlations, Bayesian analysis provided anecdotal evidence ^57^. With respect to our main analysis, we found *thermal* and *cardiac interoceptive accuracy* did not correlate with each other (*r* = 0.08, *p* = .676, *BF* = 0.239), with BF providing substantial evidence. Finally, *thermal interoceptive awareness* did not correlate with *cardiac interoceptive awareness* (*r* = −0.28, *p* = .123, *BF* = 0.685), again with BFs providing only anecdotal evidence.

### The role of thermosensitivity and interoceptive sensibility in predicting thermal interoceptive accuracy

#### Thermosensitivity and thermal interoceptive accuracy

The first linear regression analysis investigated whether performance in the thermal interoceptive task was predicted by subjective thermosensitivity (High temperature sensitivity, Solitary thermoregulation, and Social thermoregulation of the STRAQ-1 questionnaire; Heat perception and Heat warming of the ETSRS questionnaire). The results show that thermal interoceptive accuracy was predicted by Heat perception (*b*= −7.448, *t* = −2.315, *p* = .009), and Heat induced warming (*b*= 9.51, *t*= 2.79, *p* = .010) but not by High temperature sensitivity, solitary, or social thermoregulation (all *p*’s > .05). (*Figure 5*)

**Figure 5.**
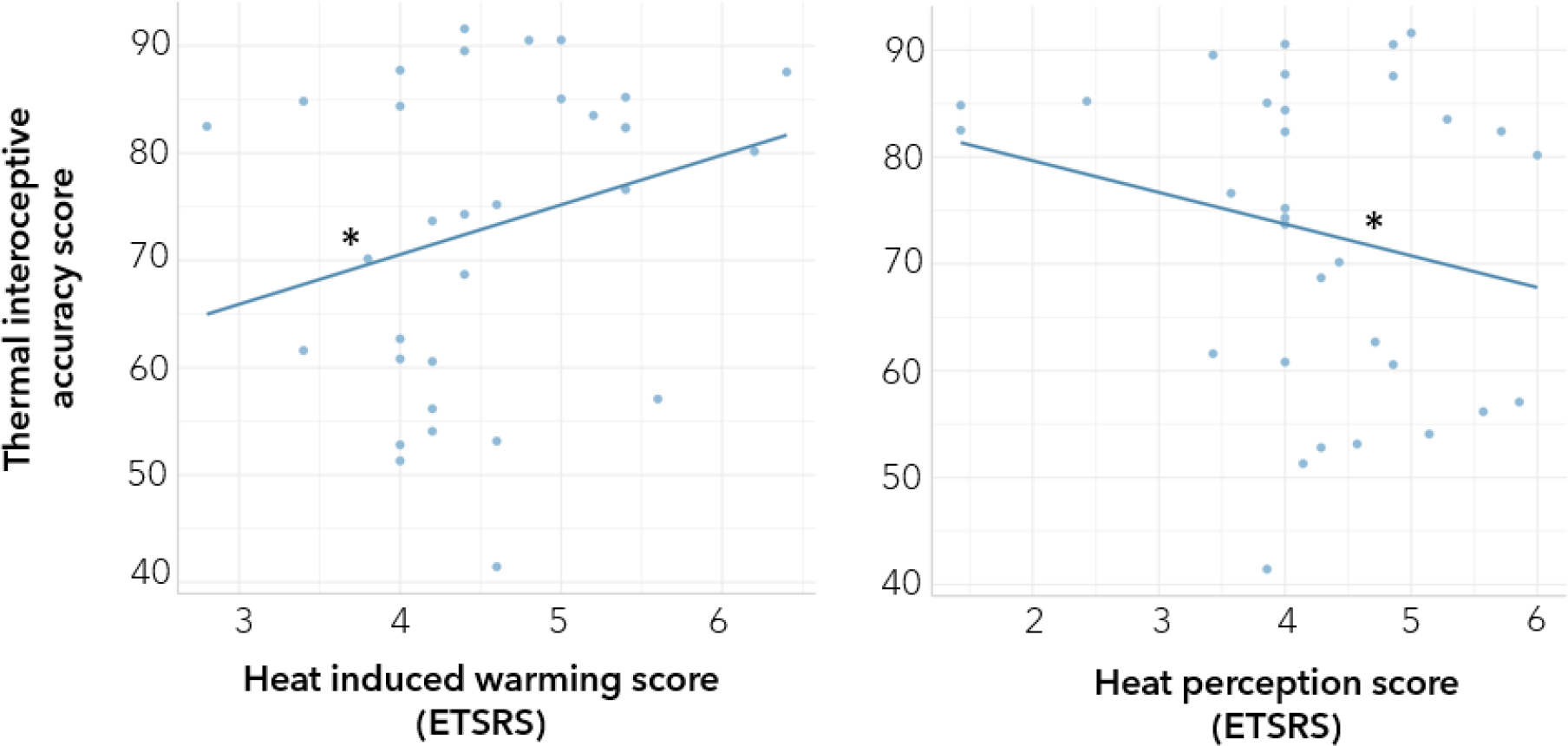
Linear relationship between thermal interoceptive accuracy and self-reported thermosensitivity. Thermal interoceptive accuracy is positively predicted by scores in Heat induced warming and negatively predicted by Heat perception. * Indicates a significant slope at p ≤ .01

#### Interoceptive sensibility and thermal interoceptive accuracy

The second linear regression analysis investigated whether task performance was predicted by subjective thermosensitivity. The results show that thermal interoceptive accuracy was predicted by Self-Regulation of the MAIA-2 (*b*=9.07, *t*= 2.41, *p* = .024). There was no significant effect of Noticing, Not-Distracting, Not-Worrying, Attention Regulation, Emotional Awareness, Body Listening, Trusting, or the BPQ-SF on thermal interoceptive accuracy (all *p*’s > .0). (*Figure 6*)

**Figure 6.**
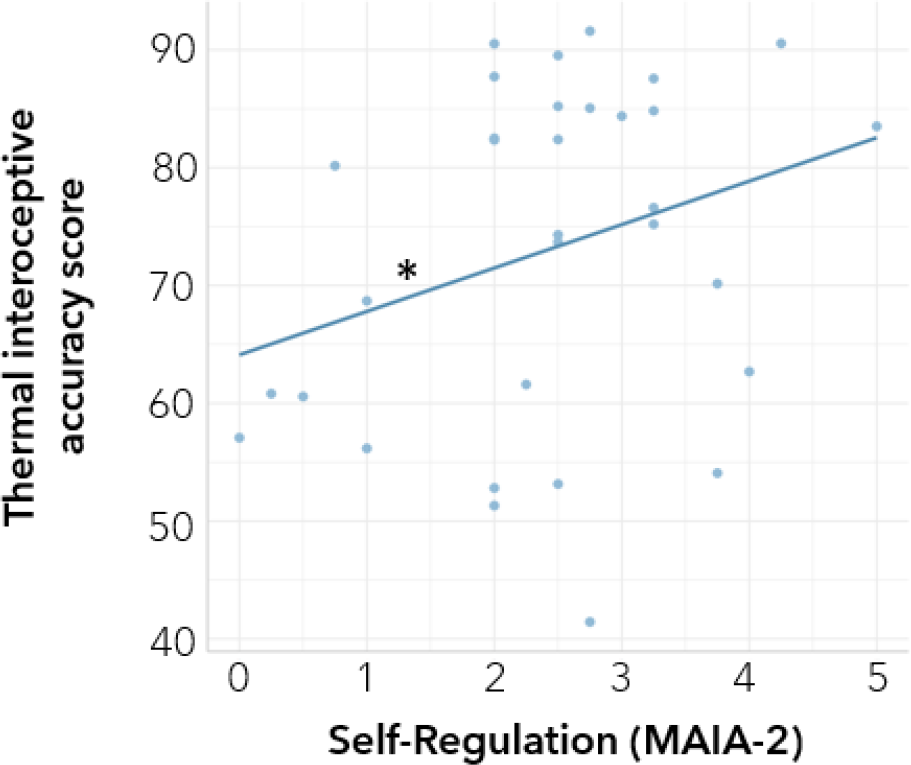
Linear relationship between thermal interoceptive accuracy and self-reported interoceptive sensibility. Self-Regulation (MAIA-2) positively predicted thermal interoceptive accuracy. * indicates a significant slope at p < .05

## Discussion

In the present study we report on the development of a task to measure awareness of changes in peripheral body temperature. In 15 trials, participants received 20-second infrared thermal stimulations to the palm of their right hand. We used five different temperature intensities (0%, 25%, 50%, 75%, and 100% of the infrared lightbulb’s maximum intensity), each presented three times. After each stimulation, participants judged how much their hand temperature had changed following the stimulation (*perceived hand temperature change*). Participants’ responses were compared to a *standardised hand temperature change*, derived from the *real hand temperature change in °C* as measured using a thermal camera. From this comparison we were able to derive a measure of *thermal interoceptive accuracy*.

### Measuring awareness of changes in peripheral body temperature

The average *real hand temperature change* was greater with each increasing stimulation intensity, evidencing that our thermal manipulation was effective in producing incremental thermal variations in the participants’ hand. Mirroring these results, participants also reported perceiving increasingly larger temperature changes (*perceived hand temperature change*) as the intensity of the stimulations increased. For what concerns our measure of *thermal interoceptive accuracy*, it showed good inter-individual variability, which suggests its potentiality as a tool for the assessment of each individual’s thermal awareness. While participants were generally better at detecting when there was no (or a very low) change in hand temperature following the baseline stimulation (0% stimulation) compared to all other stimulation conditions, there was no significant difference in their ability to accurately detect temperature changes following the other stimulation intensities (i.e., 25%, 50%, 75% and 100%). This result suggests that *thermal interoceptive accuracy* is stable across stimulation intensity. Our findings also show that *thermal interoceptive accuracy* did not correlate with *thermal interoceptive awareness*. This is in line with evidence supporting the idea that interoceptive accuracy and awareness generally rely on distinct, unrelated processes, with the exception of participants who score very high in interoceptive accuracy ^54^.

*Thermal interoceptive accuracy* was positively predicted by participants’ self-reported perception of how quickly or intensely different body parts, e.g., hands, feet, the torso, become warm when exposed to warm environments compared to other people (*Heat induced warming* from the ETSRS questionnaire) and negatively predicted by the tendency to always feel warm in different types of environments and/or situations, e.g., indoors, outdoors, in bed, when concentrating, when watching TV or reading (*Heat perception* from the ETSRS), suggesting that the task construct indeed reflects a general, and more accurate, awareness of changes in body temperature. It was not predicted by how bothered by high temperatures participants are (*High temperature sensitivity* from the STRAQ-1), which reflects a more qualitative aspect associated with experiencing warm temperatures, nor by *Solitary* or *Social behavioural thermoregulation* (from the STRAQ-1). However, thermal interoceptive accuracy was also positively predicted by the ability to self-regulate through attention to body sensations (*Self-Regulation* of the MAIA-2), highlighting a potential link between body temperature awareness and self-regulatory behaviour. Future research should better investigate the relationship between thermal awareness and the tendency to take action to maintain optimal body temperature and homeostasis. Finally, task performance was not significantly predicted by other MAIA-2 subscales (including sub-scales of basic body awareness i.e., *Noticing* or *Body Listening*) or by the *BPQ-SF*, suggesting that just as cardiac interoceptive, thermal interoceptive accuracy is not directly related to general interoceptive sensibility.

### Thermoception and cardiac interoception

The present study also shows that *thermal interoceptive accuracy* (performance in the thermoception task) did not correlate with *cardiac interoceptive accuracy* (performance in the heartbeat counting task), nor did *thermal* and *cardiac interoceptive awareness*. Similarly Crucianelli and collegues also observed the absence of a significant association between performance in a thermal matching task and that in the heartbeat counting task ^26^. Crucially, the results of these studies align with evidence consistently suggesting that conscious perception of interoceptive signals may differ across interoceptive channels ^23–25^. Very importantly, these findings should admonish against using cardiac interoception (or any measure related to a specific interoceptive domain) as a proxy for interoceptive accuracy in general and encourage researchers to rely on multiple interoceptive tests when assessing how different individuals process visceral signals, or to focus on a limited number of modalities only when driven by specific hypotheses.

We also found that the participants were more accurate in the thermoception task compared to the heartbeat counting task. It is possible that the difference between the two performances reflects the saliency, and possibly a greater awareness, of thermal signals (compared to cardiac) in daily life. Indeed, heartbeats produce faint and transient signals that we are often unaware of, as they become salient only under specific circumstances, such as physical exertion, tachycardia, and panic attacks ^58^. Thus, people are not accustomed to paying attention to their heartbeat like they are required to do during the HCT. Oppositely, the great variability of the environmental temperatures we dwell in, particularly in these global warming times, requires individuals to take direct, behavioural actions to control body temperature and preserve thermoneutrality. As such, we argue that people may be more accustomed to recognising temperature fluctuations than they are to recognising cardiac fluctuations, possibly explaining the difference in performance we observed in our two interoceptive tasks.

### Limitations

An important aspect of the thermoception task we propose here is that it is successful in inducing changes of peripheral body temperature without having a tactile confound, so that awareness of these can be assessed. However, what remains unclear is whether the processing of changes in hand temperature may rely on processes that are different from those regarding shifts in whole-body and facial temperature. Future research may attempt to disentangle whether performance in this task may relate to how people perceive changes in facial temperature, for example using heat lamps and paradigms in which participants can keep their eyes closed to avoid visual cues, and whole-body temperature, possibly by inducing changes in room temperature. Indeed, research has evidenced changes in whole-body and facial temperature changes in emotionally arousing situations ^37–41^ and in social contexts of social connection, ^42,59^ social exclusion ^43,44,60,61^, dishonest behaviour ^45,46^. We submit that if perception of temperature is related across different body areas (peripheral, facial, and whole-body), our task may provide an effective and ecological measurement of general thermosensitivity. Furthermore, while our study investigated the relationship of thermoception with cardiac interoception, more research is needed to further understand the way in which the present task relates to other measures of awareness of the physiological states of the body. Particularly, as temperature perception is often mediated by the skin, it would be important to assess the relationship of our task with other measures of touch-mediated thermosensitivity ^48,62^.

### Conclusive remarks

Conclusively, we developed a task measuring awareness of changes in peripheral body temperature (participants’ right hand) free from the tactile confound and through a safe, non-invasive stimulation procedure that did not cause excessive discomfort to participants. Hence, the thermoception task we developed appears to have the potential to become an important tool for examining awareness of the physiological state of the body in research investigating thermoregulatory behaviour in emotionally arousing contexts, where body temperature changes often reveal changes in autonomic function. Furthermore, research suggests that human social relations may be grounded in the evolutionary need for physical warmth that guides attachment formation in development, thus linking body temperature to social relations (see Ijzerman et al., 2015). Thus, tasks that specifically assess thermal interoception may be particularly relevant when investigating social situations in which social connection is at stake, like those involving the threat of negative social evaluation and thus social exclusion.

## Data availability

The anonymized datasets analyzed in the current study are available in the OSF repository, https://osf.io/6bzm4/?view_only=fbe54a3e1e4d4259bc4849fc14db1c00

## Code Availability

The scripts used for presenting the experimental stimuli and collecting participants’ responses as well as for analysis are available upon request to the corresponding author

## Ethics approval

The study was approved by the Ethics Committee of the Santa Lucia Hospital I.R.C.C.S. and was in accordance with the 1964 declaration of Helsinki.

## Author contributions

A.V., G.P., M.S.P., M.SC, and S.M.A. conceived and planned the experiment. A.V., M.SC., & M.SP. collected data. A.V. & M.SP. performed the preprocessing of the data. A.V. performed the main analysis and G.P., M.S.P., & M.SC contributed to the interpretation of the results. A.V. wrote the original draft and G.P., M.S.P., M.SC. and S.M.A. revised the manuscript. S.M.A. acquired funding and resources and S.M.A. & G.P. supervised the project. All authors read and agreed to the published version of the manuscript.

